# Deep phylo-taxono genomics reveals *Xylella* as a variant lineage of plant associated *Xanthomonas* with *Stenotrophomonas* and *Pseudoxanthomonas* as misclassified relatives

**DOI:** 10.1101/2021.08.22.457248

**Authors:** Kanika Bansal, Sanjeet Kumar, Amandeep Kaur, Anu Singh, Prabhu B. Patil

## Abstract

Genus *Xanthomonas* is a group of phytopathogens which is phylogenetically related *to Xylella, Stenotrophomonas* and *Pseudoxanthomonas* following diverse lifestyles. *Xylella* is a lethal plant pathogen with highly reduced genome, atypical GC content and is taxonomically related to these three genera. Deep phylo-taxono-genomics reveals that *Xylella* is a variant *Xanthomonas* lineage that is sandwiched between *Xanthomonas* species. Comparative studies suggest the role of unique pigment and exopolysaccharide gene clusters in the emergence of *Xanthomonas* and *Xylella* clades. Pan genome analysis identified set of unique genes associated with sub-lineages representing plant associated *Xanthomonas* clade and nosocomial origin *Stenotrophomonas*. Overall, our study reveals importance to reconcile classical phenotypic data and genomic findings in reconstituting taxonomic status of these four genera.

**Significance Statement:** *Xylella fastidiosa* is a devastating pathogen of perennial dicots such as grapes, citrus, coffee, and olives. The pathogen is transmitted by an insect vector to its specific host wherein the infection leads to complete wilting of the plants. The genome of *X. fastidiosa* is extremely reduced both in terms of size (2Mb) and GC content (50%) when compared with its relatives such as *Xanthomonas, Stenotrophomonas, and Pseudoxanthomonas* that have higher GC content (65%) and larger genomes (5Mb). In this study, using systematic and in-depth genome-based taxonomic and phylogenetic criteria along with comparative studies, we assert the need of unification of *Xanthomonas* with its misclassified relatives (*Xylella*, *Stenotrophomonas* and *Pseudoxanthomonas*). Interestingly, *Xylella* revealed itself as a minor lineage embedded within two major *Xanthomonas* lineages comprising member species of different hosts.

## Introduction

Family *Lysobacteraceae* (*Xanthomonadaceae*) (CHRISTENSEN & Cook, 1978; S. Naushad et al., 2015) harbors bacterium of diverse ecological niches. Within this family, *Xanthomonas*, *Stenotrophomonas, Xylella* and *Pseudoxanthomonas* are closely related genera which forms a phylogroup (referred to as XSXP phylogroup in this study) (Kumar, Bansal, Patil, & Patil, 2019). *Xanthomonas, Xylella* and *Pseudoxanthomonas* are characterized by a yellow pigment xanthomonadin and exopolysaccharide xanthan gum production (Biswas, Chakraborty, Sarkar, & Naidu, 2017; da Silva, Vettore, Kemper, Leite, & Arruda, 2001; He, Cao, & Poplawsky, 2020; Katzen et al., 1998; Lu et al., 2008; Rajagopal, Sundari, Balasubramanian, & Sonti, 1997). XSXP phylogroup have a long history of taxonomic reshuffling based on phenotypic, morphological characteristics and genotypic methods. These genotypic methods were based on single gene such as 16S rRNA, *rpoB* or *gyrB* gene or multiple housekeeping genes (Parkinson, Cowie, Heeney, & Stead, 2009; Saddler & Bradbury, 2005; Yarza et al., 2010; Yilmaz et al., 2014). Taxonomic and phylogenetic relationship amongst XSXP based on classical methods have been always debatable. For instance, basionym of *Xanthomonas* and *Stenotrophomonas* were *Bacillus campestris* by Pammel in 1895 (Pammel, 2017) and *Pseudomonas maltophilia* respectively (Hugh & Ryschenkow, 1961). Further, *Bacillus campestris* including few groups of the plant pathogens were later classified as *Xanthomonas campestris* (Dowson, 1939). Similarly, *Pseudomonas maltophilia* was transferred to genus *Xanthomonas* as *X. maltophilia* (Swings, De Vos, den MOOTER, & De Ley, 1983) and later it was designated as a distinct and new genus *Stenotrophomonas* (Palleroni & Bradbury, 1993). Similarly, *Xylella* (Wells et al., 1987) and *Pseudoxanthomonas* (Finkmann, Altendorf, Stackebrandt, & Lipski, 2000) were defined as species related to *Xanthomonas*. However, these taxonomic assignments remain inconclusive as these were based on few evidences of classical taxonomy such as 16S rRNA identity or biochemical properties etc. Current standing in nomenclature of *Lysobacteraceae* (https://lpsn.dsmz.de/family/lysobacteraceae) classifies XSXP phylogroup as four distinct genera (i.e., *Xanthomonas, Pseudoxanthomonas, Stenotrophomonas* and *Xylella*). This is based on conserved sequence indels (CSI) and phylogeny of only 28 proteins (H. S. Naushad & Gupta, 2013; S. Naushad et al., 2015). However, CSI phylogeny also suggests intermingling of some species of these four genera (H. S. Naushad & Gupta, 2013; S. Naushad et al., 2015). Further, genome taxonomy database (GTDB) also supports inclusion of several genera in the XSXP phylogroup. GTDB classification is based on 120 conserved genes and relative evolutionary divergence values. However, incongruence to the GTDB proposed taxonomy is reported in some previous focused studies (Liao, Lin, Li, Qu, & Tian, 2020; Zheng et al., 2020). GTDB proposed classification for XSXP phylogroup is not yet reported in literature and hence, its evolutionary and taxonomic status is worth discussing.

Taxonomy and phylogeny go hand in hand. Amongst XSXP, existence of two major phylogroups within genus *Xanthomonas* (group 1 and group 2) appears as early as in 1960s (Colwell & Liston, 1961). Later on, single gene phylogeny of 16S rRNA and *gyrB* (Hauben, Vauterin, Swings, & Moore, 1997; Parkinson et al., 2007; Pieretti et al., 2009; Studholme et al., 2011) defined the group 1 as early branching. Group 1 is highly diverse which constitutes species like *X. albilineans* with reduced genome (Pieretti et al., 2009)*, X. sontii* following non-pathogenic lifestyle (Bansal, Kaur, et al., 2019; Bansal, Kumar, & Patil, 2020; Bansal, Midha, et al., 2019a) in addition to several pathogens like *X. sacchari, X. translucens,* etc. Whereas, group 2 is the largest and constitutes well-described pathogens such as *X. oryzae, X. citri, X. campestris* etc. (Hauben et al., 1997; Parkinson et al., 2007).

However, phylogeny of two devastating phytopathogenic genera *Xanthomonas* and *Xylella* on the basis of single or multi-locus housekeeping genes have been debatable over the years (Pieretti et al., 2009; Ryan et al., 2011; Studholme et al., 2011). These studies did not resolve their relationship and either places *Xylella* within group 1 and group 2 (Pieretti et al., 2009) of *Xanthomonas* or as a distinct monophyletic lineage (Ryan et al., 2011). Furthermore, 16S rRNA based phylogeny have also placed *Xanthomonas* and *Stenotrophomonas* as coherent group excluding *Xylella* (Pieretti et al., 2009).

With recent advancements, genomics is at the center of revolution in bacterial taxonomy and phylogeny. Whole genome comparisons of *Xanthomonas* and *Xylella* have revealed high degree of identity and co-linearity of their chromosome backbone (Monteiro-Vitorello et al., 2005; Van Sluys et al., 2002). Even though, their genomes have diverged by potential indels mediated by mobile genetic elements, *Xylella* shares 74% of the genes with *Xanthomonas* (Moreira et al., 2004) (Monteiro-Vitorello et al., 2005). *Xylella* genome size is roughly two-third of the *Xanthomonas* genome with minimal complement of genes required for its survival within host (Lu et al., 2008). Like *Xylella*, *Xanthomonas albilineans* is also a xylem-limited plant pathogen with reduced genome and characterized by the absence of Hrp-T3SS (hypersensitive response and pathogenicity–type III secretion system (Pieretti et al., 2009). However, genome reduction events in genus *Xanthomonas* are not limited to group 2 (*Xanthomonas albilineans*) but also reported in group 1 i.e., *Xanthomonas vasicola* (Rodriguez-R et al., 2012). All these preliminary genome level investigations based on limited strains provide certain clues of *Xanthomonas* and *Xylella* relatedness. Yet small genome size, low GC content and dual lifestyle of *Xylella* has compelled taxonomists to consider it as a different genus (S. Chatterjee, R. P. Almeida, & S. Lindow, 2008a). However, in microbial world genome erosion cannot warranty for a new genus rather it leads to a highly specialized pathogen. For instance, *Mycobacterium leprae* having almost 50% of the genome reduced as compared to *M. tuberculosis* (Gómez-Valero, Rocha, Latorre, & Silva, 2007).

Phylogenetic investigation which led to current standing in nomenclature was based on just 28 conserved proteins and that too of less number of species of XSXP phylogroup. Moreover, intermingled phylogeny of *Xanthomonas* and *Xylella* in the CSI phylogeny was also overlooked in the current standing in nomenclature (https://lpsn.dsmz.de/family/lysobacteraceae) (H. S. Naushad & Gupta, 2013; S. Naushad et al., 2015). Hence, this does not provide true evidence for considering XSXP phylogroup as four distinct genera. The increasing evidences of relatedness amongst the XSXP phylogroups beyond the genus level is gradually strengthening. However, to get robust taxonomy on the basis of phylogenomic framework a comprehensive genome level investigation considering all representative species of XSXP is required. Present deep whole genome based phylogenetic and evolutionary investigation reveals that all the four genera indeed belong to the same genus *Xanthomonas*. Such a deep phylo-taxono-genomic or DEEPT genomic study of XSXP have major implications in understanding the co-evolution of microbes with their host plants.

## Results

### Genomic features of XSXP phylogroup

Genomic features of type strains of the XSXP phylogroup are summarized in table 1. Genome size of *Xanthomonas* is around 5 Mb and around 3-5 Mb for *Pseudoxanthomonas* and *Stenotrophomonas,* whereas, genome size of *Xylella* is around 2.5 Mb. This reduction in genome size is also reflected in number of coding sequences that is in the range of 3000 to 4000 for *Xanthomonas, Stenotrophomonas* and *Pseudoxanthomonas,* but around 2100 for *Xylella.* Furthermore, average GC content of the genome for all the members of *Xanthomonas*, *Stenotrophomonas* and *Pseudoxanthomonas* is ~65% except for *Xylella* that has a GC content of around 51%. In spite of dramatic genome reduction *Xylella* is displaying comparable number of tRNAs i.e., 48.

**Table 1:** Metadata of the strains used in the present study. Following data was obtained from list of prokaryotic names with standing in nomenclature (http://www.bacterio.net/) and NCBI (https://www.ncbi.nlm.nih.gov/). Star marked columns are obtained in the present study. The data which is not available is marked with - across Host taxonomy, Lifestyle, Location and Isolation year columns.

| Strain | Genome size | CDS | Genome status | Source/Host | Host taxonomy | Lifestyle | Location | Isolation year | GC% | # tRNA | Completeness*/Contaminations* |
| --- | --- | --- | --- | --- | --- | --- | --- | --- | --- | --- | --- |
| *Xanthomonas vasicola* NCPPB 2417<sup>T</sup> | 4.9 | 3960 | Draft | *Sorghum vulgare* | Tracheophytes, Angiosperms, Monocots, Commelinids, Poales, Poaceae | Pathogenic | New Zealand | 1969 | 63.3 | 51 | 100/0.07 |
| *Xanthomonas oryzae* ATCC 35933<sup>T</sup> | 4.2 | 3385 | Draft | *Oryza sativa* (leaf blight and wilt) | Tracheophytes, Angiosperms, Monocots, Commelinids, Poales, Poaceae | Pathogenic | - | - | 63.9 | 49 | 100/0 |
| *Xanthomonas euvesicatoria* 85-10 | 5.1 | 4416 | Complete | Infected pepper plant | Tracheophytes, Angiosperms, Eudicots, Asterids, Solanales, Solanaceae | Pathogenic | - | - | 64.7 | 54 | 99.64/0 |
| *Xanthomonas alfalfae* LMG 495<sup>T</sup> | 5.0 | 3937 | Draft | *Medicago sativa* | Tracheophytes, Angiosperms, Eudicots, Rosids, Fabales, Fabaceae | Pathogenic | Belgium: Merelbeke | 2014 | 64.8 | 44 | 98.79/2.99 |
| *Xanthomonas perforans* CFBP 7293<sup>T</sup> | 5.0 | 4206 | Draft | Leaves of *Solanum lycopersicum* | Tracheophytes, Angiosperms, Eudicots, Asterids, Solanales, Solanaceae | Pathogenic | USA: Florida | - | 64.9 | 54 | 99.64/0.2 |
| *Xanthomonas phaseoli* CFBP 412 | 5.0 | 4175 | Complete | *Phaseolus vulgaris* | Tracheophytes, Angiosperms, Eudicots, Rosids, Fabales, Fabaceae | Pathogenic | USA | - | 65 | 54 | 99.64/0.24 |
| *Xanthomonas citri* LMG 9322<sup>T</sup> | 5.1 | 4355 | Draft | *Citrus aurantifolia* | Tracheophytes, Angiosperms, Eudicots, Rosids, Sapindales, Rutaceae | Pathogenic | USA: Florida | 1915 | 64.6 | 50 | 98.19/1.09 |
| *Xanthomonas axonopodis* DSM 3585<sup>T</sup> | 4.4 | 3263 | Draft | *Axonopus scoparius* | Tracheophytes, Angiosperms, Monocots, Commelinids, Poales, Poaceae | Pathogenic | Colombia | 1949 | 64.5 | 50 | 99.52/0.12 |
| *Xanthomonas prunicola* CFBP 8353<sup>T</sup> | 5.3 | 4208 | Draft | *Prunus persica* var. nectarina | Tracheophytes, Angiosperms, Eudicots, Rosids, Rosales, Rosaceae | Pathogenic | Spain: Murcia, Abaran | 2015 | 64 | 53 | 100/0 |
| *Xanthomonas bromi* CFBP 1976<sup>T</sup> | 4.9 | 3845 | Draft | *Bromus carinatus* | Tracheophytes, Angiosperms, Monocots, Commelinids, Poales, Poaceae | Pathogenic | France | 1980 | 64 | 53 | 100/0 |
| *Xanthomonas nasturtii* WHRI 8853<sup>T</sup> | 4.8 | 3981 | Draft | Watercress | Tracheophytes, Angiosperms, Eudicots, Rosids, Brassicales, Brassicaceae | Pathogenic | USA: Florida | 2014 | 64.5 | 56 | 100/0.36 |
| *Xanthomonas cynarae* CFBP 4188<sup>T</sup> | 5.0 | 4102 | Draft | *Cynara scolymus* | Tracheophytes, Angiosperms, Eudicots, Asterids, Asterales, Asteraceae | Pathogenic | France: Bretagne | 1996 | 63.7 | 51 | 100/0.41 |
| *Xanthomonas gardneri* ICMP 7383 | 5.3 | 4523 | Complete | Tomato field | Tracheophytes, Angiosperms, Eudicots, Asterids, Solanales, Solanaceae | Pathogenic | New Zealand | 1980 | 63.5 | 53 | 99.64/0.68 |
| *Xanthomonas hortorum* WHRI 7744<sup>T</sup> | 5.5 | 4427 | Draft | *Hedera helix* | Tracheophytes, Angiosperms, Eudicots, Asterids, Apiales, Araliaceae | Pathogenic | USA | 1961 | 63.6 | 54 | 100/1.03 |
| *Xanthomonas populi* CFBP 1817<sup>T</sup> | 4.3 | 3222 | Draft | *Populus x canadensis* cv. regenerata) | Tracheophytes, Angiosperms, Eudicots, Rosids, Malpighiales, Salicaceae | Pathogenic | Oise, Noyon | 1957 | 63.3 | 48 | 99.64/0.14 |
| *Xanthomonas arboricola* CFBP 2528<sup>T</sup> | 5.0 | 4124 | Draft | Twig from a walnut tree | Tracheophytes, Angiosperms, Eudicots, Asterids, Fagales, Juglandaceae | Pathogenic | New Zealand | 1956 | 65.5 | 51 | 100/0.41 |
| *Xanthomonas fragariae* PD 885 | 4.2 | 3263 | Complete | Diseased strawberry | Tracheophytes, Angiosperms, Eudicots, Rosids, Rosales, Rosaceae | Pathogenic | - | - | 62.3 | 52 | 99.64/0.36 |
| *Xanthomonas pisi* DSM 18956 | 4.2 | 3385 | Draft | *Pisum sativum*; Leaf spot | Tracheophytes, Angiosperms, Eudicots, Rosids, Fabales, Fabaceae | Pathogenic | Japan | 1997 | 64.7 | 43 | 80.23/0 |
| *Xanthomonas dyei* CFBP 7245<sup>T</sup> | 5.3 | 4278 | Draft | *Metrosideros excelsa*;<br>Angular leaf lesion | Tracheophytes, Angiosperms, Eudicots, Rosids, Myrtales, Myrtaceae | Pathogenic | Bay of Plenty<br>Omahanui | 1993 | 64.3 | 51 | 100/0.36 |
| *Xanthomonas vesicatoria* LMG 911<sup>T</sup> | 5.1 | 4343 | Complete | *Lycopersicon esculentum* | Tracheophytes, Angiosperms, Eudicots, Asterids, Solanales, Solanaceae | Pathogenic | New Zealand | 1955 | 64.3 | 55 | 100/0.36 |
| *Xanthomonas melonis* CFBP 4644 | 4.6 | 3872 | Draft | *Cucumis melo* | Tracheophytes, Angiosperms, Eudicots, Rosids, Cucurbitales, Cucurbitaceae | Pathogenic | Brazil | 1974 | 66.1 | 52 | 99.64/0.18 |
| *Xanthomonas cucurbitae* CFBP 2542<sup>T</sup> | 4.4 | 3553 | Draft | *Cucurbita maxima* | Tracheophytes, Angiosperms, Eudicots, Rosids, Cucurbitales, Cucurbitaceae | Pathogenic | New Zealand | 1968 | 65.4 | 51 | 100/0.36 |
| *Xanthomonas floridensis* WHRI 8848<sup>T</sup> | 5.2 | 4286 | Draft | Watercress | Tracheophytes, Angiosperms, Eudicots, Rosids, Brassicales, Brassicaceae | Pathogenic | USA: Florida | 2014 | 65.4 | 52 | 100/0.36 |
| *Xanthomonas cassavae* CFBP 4642<sup>T</sup> | 5.2 | 4218 | Draft | *Manihot esculenta* | Tracheophytes, Angiosperms, Eudicots, Rosids, Malpighiales, Euphorbiaceae | Pathogenic | Malawi | 1951 | 65.2 | 51 | 99.64/0.93 |
| *Xanthomonas codiae* CFBP 4690 | 5.0 | 4125 | Draft | *Codiaeum variegatum* | Tracheophytes, Angiosperms, Eudicots, Rosids, Malpighiales, Euphorbiaceae | Pathogenic | USA: Florida | 1987 | 66 | 53 | 99.28/0 |
| *Xanthomonas campestris* pv. *campestris* ATCC 33913<sup>T</sup> | 5.0 | 4179 | Complete | Plant (*Brassica oleracea* var. *gemmifera*) | Tracheophytes, Angiosperms, Eudicots, Rosids, Brassicales, Brassicaceae | Pathogenic | - | - | 65.1 | 53 | 99.64/0 |
| *Xanthomonas maliensis* LMG 27592<sup>T</sup> | 5.2 | 4197 | Draft | *Oryza sativa* | Tracheophytes, Angiosperms, Monocots, Commelinids, Poales, Poaceae | Non-pathogenic | Mali: Africa | 2009 | 66.2 | 53 | 96.74/0.3 |
| *Xylella taiwanensis* PLS229<sup>T</sup> | 2.7 | 2047 | Draft | Pear | Tracheophytes, Angiosperms, Eudicots, Rosids, Rosales, Rosaceae | Pathogenic | Taiwan | - | 53 | 47 | 89.88/0.12 |
| *Xylella fastidiosa* ATCC 35879<sup>T</sup> | 2.5 | 2105 | Draft | Grapevine | Tracheophytes, Angiosperms, Eudicots, Rosids, Vitales, Vitaceae | Pathogenic | USA: Florida | 1987 | 51.8 | 49 | 99.23/0 |
| *Stenotrophomonas panacihumi* JCM 16536<sup>T</sup> | 3.9 | 3497 | Draft | Soil of a ginseng field | - | Environmental | South korea | 2010 | 68.8 | 51 | 97.59/0.34 |
| *Xanthomonas sontii* PPL1<sup>T</sup> | 4.8 | 3937 | Draft | *Oryza sativa* | Tracheophytes, Angiosperms, Monocots, Commelinids, Poales, Poaceae | Non-pathogenic | India | 2012 | 69 | 51 | 96.31/0.22 |
| *Xanthomonas sacchari* CFBP 4641<sup>T</sup> | 4.9 | 4098 | Draft | *Saccharum officinarum* | Tracheophytes, Angiosperms, Monocots, Commelinids, Poales, Poaceae | Pathogenic | Guadeloupe | 1980 | 69.1 | 51 | 100/0.41 |
| *Xanthomonas albilineans* CFBP 2523<sup>T</sup> | 3.6 | 2967 | Draft | *Saccharum officinarum* | Tracheophytes, Angiosperms, Monocots, Commelinids, Poales, Poaceae | Pathogenic | Fiji | 1961 | 63.1 | 49 | 99.96/0.34 |
| *Xanthomonas hyacinthi* CFBP 1156 | 4.9 | 4003 | Draft | *Hyacinthus orientalis* | Tracheophytes, Angiosperms, Monocots, Asparagales, Asparagaceae | Pathogenic | Netherlands | 1958 | 68.1 | 51 | 100/0.03 |
| *Xanthomonas translucens* DSM 18974<sup>T</sup> | 4.4 | 3788 | Draft | *Hordeum vulgare* | Tracheophytes, Angiosperms, Monocots, Commelinids, Poales, Poaceae | Pathogenic | - | - | 67.7 | 53 | 98.5/0.12 |
| *Xanthomonas theicola* CFBP 4691 | 4.5 | 3630 | Draft | *Camellia sinensis* | Tracheophytes, Angiosperms, Eudicots, Asterids, Ericales, Theaceae | Pathogenic | Japan | 1974 | 68.4 | 49 | 99.95/0 |
| *Stenotrophomonas ginsengisoli* DSM 24757<sup>T</sup> | 3.3 | 3001 | Draft | Soil from ginseng field | - | Environmental | South Korea | 2010 | 65.9 | 76 | 98.72/0.76 |
| *Stenotrophomonas koreensis* DSM 17805<sup>T</sup> | 3.0 | 2690 | Draft | Compost | - | -- | South Korea | 2003 | 66.1 | 75 | 98.16/0.79 |
| *Stenotrophomonas terrae* DSM 18941<sup>T</sup> | 4.4 | 3730 | Draft | Soil | - | Environmental | Belgium | 2007 | 63.9 | 60 | 99.19/1.38 |
| *Stenotrophomonas humi* DSM 18929<sup>T</sup> | 4.1 | 3591 | Draft | Soil | - | Environmental | Belgium | 2007 | 64 | 62 | 99.6/1.03 |
| *Stenotrophomonas pictorum* JCM 9942<sup>T</sup> | 3.3 | 3150 | Draft | Soil | - | Environmental | - | 1928 | 66 | 61 | 99.84/0.32 |
| *Stenotrophomonas nitritireducens* DSM 12575<sup>T</sup> | 3.9 | 3476 | Draft | Laboratory scale biofilters supplied with ammonia or dimethyl disulfide and ammonia | - | - | Germany | 1997 | 68.3 | 54 | 95.5/0.78 |
| *Stenotrophomonas acidaminiphila* JCM 13310<sup>T</sup> | 3.9 | 3511 | Draft | Sludge from a lab-scale anaerobic chemical waste water reactor | - | Environmental | Mexico | 1999 | 68.8 | 56 | 96.69/1.51 |
| *Stenotrophomonas daejeonensis* JCM 16244<sup>T</sup> | 3.2 | 2888 | Draft | Sewage water | - | Environmental | South Korea | 2010 | 68.6 | 56 | 99.59/0.86 |
| *stenotrophomonas hibiscicola* ATCC 19867<sup>T</sup> | 4.4 | 3928 | Draft | Plant | - | - | - | - | 66.4 | 64 | 100/0 |
| *Stenotrophomonas pavanii* DSM 25135<sup>T</sup> | 4.3 | 3783 | Draft | Stems of sugar cane | Tracheophytes, Angiosperms, Monocots, Commelinids, Poales, Poaceae | Symbiotic | Brazil | 2011 | 66.2 | 67 | 100/0 |
| *Stenotrophomonas africana* LMG 22072<sup>T</sup> | 4.5 | 3991 | Draft | Cerebrospinal fluid | - | Opportunistic pathogen | Democratic Republic of the Congo | 1994 | 66.3 | 65 | 97.05/0.23 |
| *Pseudomonas geniculata* JCM 13324<sup>T</sup> | 4.8 | 4339 | Draft | Tap water | - | Environmental | USA | 1895 | 66.2 | 66 | 99.76/0.41 |
| *Stenotrophomonas maltophilia* NCTC 10257<sup>T</sup> | 5.0 | 4491 | Complete | Mouth | - | Opportunistic pathogen | Germany | 1900/1961 | 66.1 | 73 | 100/0.34 |
| *Stenotrophomonas beteli* LMG 00978<sup>T</sup> | 4.4 | 3907 | Draft | *Piper betle* | Tracheophytes, Angiosperms, Magnoliids, Piperales, Piperaceae | - | Sri Lanka | 1928 | 66.8 | 67 | 99.08/0 |
| *Stenotrophomonas indicatrix* WS40<sup>T</sup> | 4.5 | 4067 | Draft | Dirty dishes | - | Environmental | Germany: Bonn | 2013 | 66.4 | 65 | 99.91/0.57 |
| *Stenotrophomonas lactitubi* M15<sup>T</sup> | 4.8 | 4378 | Draft | Milking machine biofilm | - | - | Germany: Koenigswinter-Vinzel | 2014 | 65.9 | 66 | 100/0.41 |
| *Stenotrophomonas chelatiphaga* DSM 21508<sup>T</sup> | 3.9 | 3420 | Draft | Municipal sewage sludge | - | Environmental | Russia | 2009 | 66.9 | 62 | 98.75/0.04 |
| *Stenotrophomonas bentonitica* VV6 | 4.3 | 3805 | Draft | Arctic sea water near B02 station | - | Environmental | - | - | 66.5 | 60 | 99.91/1.93 |
| *Stenotrophomonas rhizophila* DSM 14405<sup>T</sup> | 4.6 | 3971 | Draft | Root | - | Symbiotic | Germany: Rostock | - | 67.3 | 67 | 100/0.21 |
| *Pseudoxanthomonas composti* GSS 15<sup>T</sup> | 4.3 | 3664 | Draft | Compost | - | Environmental | China | - | 68.2 | 52 | 99.31/0.8 |
| *Pseudoxanthomonas spadix* DSM 18855<sup>T</sup> | 3.3 | 3042 | Draft | Oil-contaminated soil | - | Environmental | Taiwan | Before 2006 | 67.8 | 48 | 98.97/0.59 |
| *Pseudoxanthomonas daejeonensis* DSM 17801<sup>T</sup> | 3.5 | 3143 | Draft | Soil from a ginseng field | - | Environmental | - | - | 68.8 | 58 | 99.66/0.11 |
| *Pseudoxanthomonas kaohsiungensis* DSM 17583<sup>T</sup> | 3.7 | 3420 | Draft | Oil-polluted site | - | Environmental | Taiwan | 2003 | 69.7 | 52 | 99.66/1.42 |
| *Pseudoxanthomonas koreensis* KCTC 12208<sup>T</sup> | 3.0 | 2681 | Draft | Soil from a ginseng field | - | Environmental | South Korea | - | 70.1 | 52 | 99.48/0.04 |
| *Pseudoxanthomonas jiangsuensis* DSM 22398<sup>T</sup> | 3.7 | 3389 | Draft | DDT-contaminated soil | - | Environmental | China | 2008 | 70.3 | 50 | 99.31/1.03 |
| *Pseudoxanthomonas broegbernensis* DSM 12573<sup>T</sup> | 3.5 | 3024 | Draft | Ammonia-supplied biofilters | - | Environmental | Germany | Before 1998 | 70.6 | 54 | 99.66/1.74 |
| *Pseudoxanthomonas suwonensis* DSM 17175<sup>T</sup> | 3.4 | 3071 | Draft | Cotton waste composts | - | Environmental | South Korea | Before 2005 | 70.3 | 55 | 99.66/0.34 |
| *Pseudoxanthomonas taiwanensis* DSM 22914<sup>T</sup> | 3.0 | 2729 | Draft | Hot spring | - | Environmental | Taiwan | Before 2009 | 72.08 | 56 | 99.16/0.34 |
| *Pseudoxanthomonas sangjuensis* DSM 28345<sup>T</sup> | 3.2 | 2908 | Draft | Greenhouse soil | - | Environmental | Republic of Korea | Before 2014 | 68.6 | 51 | 99.95/0.46 |
| *Pseudoxanthomonas kalamensis* DSM 18571<sup>T</sup> | 3.0 | 2697 | Draft | Soil contaminated with polycyclic aromatic hydrocarbons & polychlorinated biphenyls | - | Environmental | United States of America | Before 2006 | 65.9 | 48 | 99.84/1.5 |
| *Pseudoxanthomonas dokdonensis* DSM 21858<sup>T</sup> | 3.5 | 3153 | Draft | Soil | - | Environmental | South Korea | Before 2008 | 64.4 | 50 | 99.59/0.41 |
| *Pseudoxanthomonas wuyuanensis* DSM 100640<sup>T</sup> | 4.6 | 4003 | Draft | Saline-alkaline soil | - | Environmental | China | Before 2015 | 65.7 | 50 | 100/1.08 |
| *Pseudoxanthomonas mexicana* DSM 17121<sup>T</sup> | 3.9 | 3660 | Draft | Anaerobic digester | - | Environmental | Mexico | Before 2005 | 67.4 | 54 | 99.49/1.71 |
| *Pseudoxanthomonas arseniciresistens* KAS 5_3<sup>T</sup> | 3.9 | 4049 | Draft | Arsenic contaminated ground water | - | Environmental | India | - | 66.5 | 52 | 99.95/1.19 |
| *Pseudoxanthomonas japonensis* DSM 17109<sup>T</sup> | 4.0 | 3673 | draft | Urban riverside soil |  | Environmental | Japan | Before 2005 | 67.3 | 50 | 99.95/0.76 |
| *Pseudoxanthomonas indica* P15<sup>T</sup> | 3.9 | 3593 | Draft | Open hexachlorocyclohexane dumpsite soil | - | Environmental | India | - | 65.4 | 49 | 99.89/0.47 |
| *Pseudoxanthomonas yeongjuensis* DSM 18204<sup>T</sup> | 3.9 | 3389 | Draft | Soil of a ginseng field | - | Environmental | Republic of Korea | 2005 | 65.1 | 53 | 99.95/0.76 |
| *Pseudoxanthomonas gei* KCTC 32298<sup>T</sup> | 3.4 | 3122 | Draft | Stem of *Geum aleppicum* Jacq | Tracheophytes, Angiosperms, Eudicots, Rosids, Rosales, Rosaceae | Endophyte | Taibai Mountain in Shaanxi Province, north-west China | - | 65.4 | 45 | 99.95/0.41 |
| *Pseudoxanthomonas sacheonensis*<br>DSM 19373<sup>T</sup> | 4.0 | 3582 | Draft | BTEX-contaminated soil | - | Environmental | Republic of<br>Korea | Before<br>2007 | 64.3 | 50 | 100/0.91 |

### Phylogenomics of XSXP phylogroup

To investigate the relationship amongst the member species of XSXP phylogroup, we constructed and compared genome-based phylogeny by three different approaches. Based on an earlier study of family with basonym *Xanthomonadaceae* (*Lysobacteraceae*) and order with basonym *Xanthomonadales* (*Lysobacterales)*, we used *Luteimonas mephitis* DSM12574^T^ as an outgroup of XSXP (Kumar et al., 2019). Phylogeny constructed (including XSXP phylogroup and *Luteimonas mephitis* DSM12574^T^ as outgroup) on the basis of large set of genes core to the bacterial world (>400 genes) (figure 1a) and set of genes core to XSXP phylogroup by roary (382 genes across 99-100% isolates) (Page et al., 2015) and PIRATE (1149 genes across 95-100% isolates) (Bayliss, Thorpe, Coyle, Sheppard, & Feil, 2019) (figure 1b, supplementary figure 1) correlated with each other (pl. see methods). Core genome level phylogeny revealed that among XSXP phylogroup, *Pseudoxanthomonas* members were more diverse and ancestral to other three genera. Interestingly, other three genera formed two mega species groups (MSG) i.e. plant pathogens *Xanthomonas* and *Xylella* comprised one MSG referred as XX-MSG and *Stenotrophomonas* formed another MSG referred as S-MSG. XX-MSG consist of three clades. Clade I comprises of at least 27 species that are primarily pathogens of dicot plants which was earlier considered as group 2 (Hauben et al., 1997; Parkinson et al., 2007), clade III comprises of at least 6 species that are primarily pathogen of monocot plants including *X. albilineans* with reduced genome and *X. sontii* with non-pathogenic lifestyle earlier considered as group 1. Clade II comprised both the species of *Xylella* sandwiched in between clade I and clade III of genus *Xanthomonas.* Members of clade I are pathogens of dicots, except 5 species which have monocot hosts i.e. *X. vasicola, X. oryzae, X. axonopodis, X. bromi* and *X. maliensis*. On the other hand, members of clade III are pathogens or associated with monocots, except *X. theicola* infecting dicots. Interestingly, *Stenotrophomonas panacihumi,* an environmental species, reflected as singleton phylogenetic outlier of clade I and clade II. In case of S-MSG, we found clade IV comprising of *Stenotrophomonas maltophilia* complex (Smc) of clinical origin

**Figure 1:**
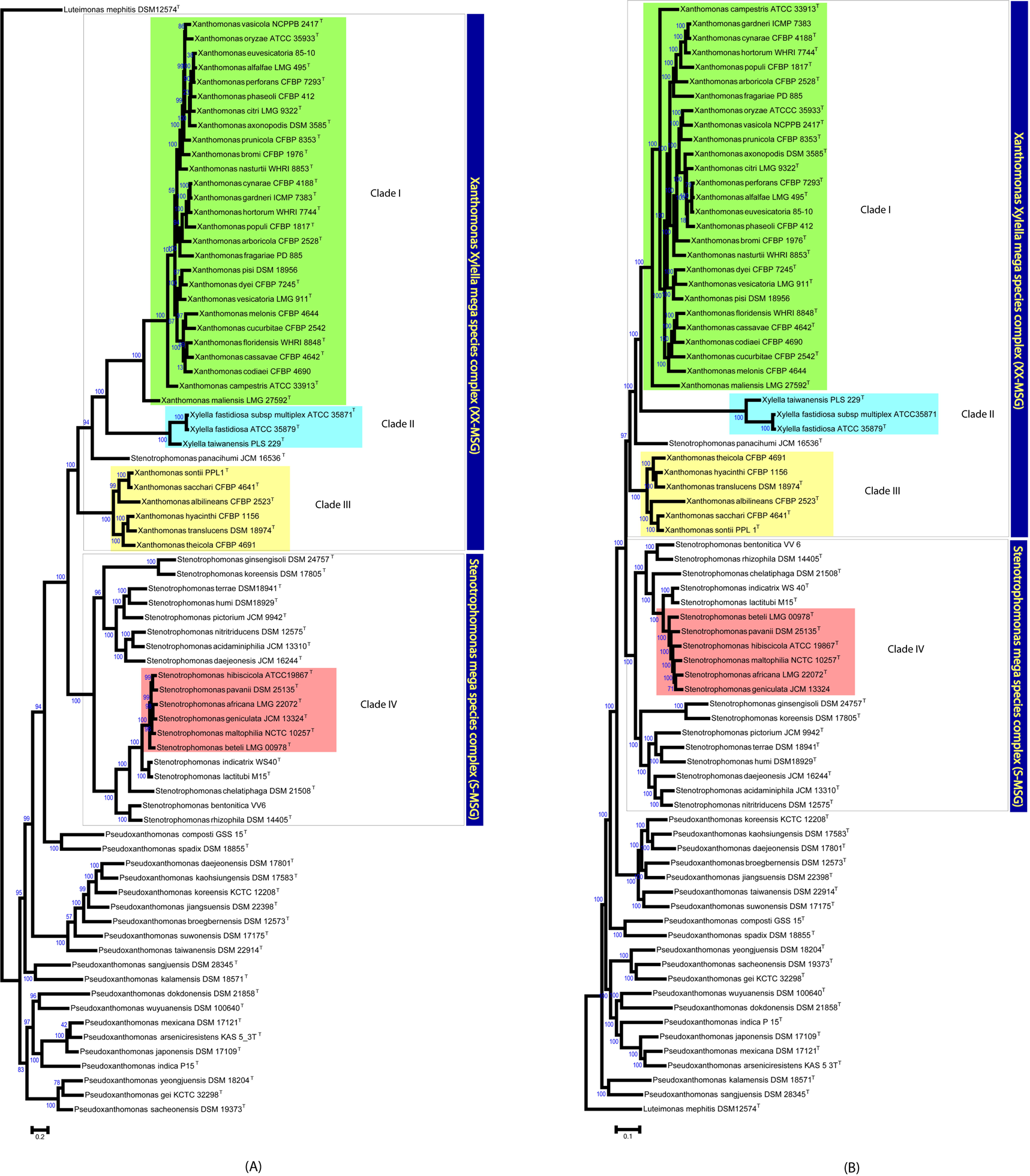
Phylogenetic construction for XSXP phylogroup. **a.** Both the mega species groups are represented by blue boxes and four species complexes are represented by colored boxes. *Luteimonas mephitis* DSM12574^T^ was used as an outgroup and bootstrap values are mentioned on the nodes. **b.** Both the mega species groups are represented by blue boxes and four species complexes are represented by colored boxes. *Luteimonas mephitis* DSM12574^T^ was used as an outgroup and bootstrap vales are mentioned on the nodes.

### Taxonogenomics of XSXP phylogroup

In order to delineate members of XSXP at genera and species level, we further carried out taxonogenomic analysis. Apart from the phylogenetic trees based on large sets of genomic markers and core genome, new criteria are also becoming available for delineating members at the genus level. For genus demarcation 73.98% average nucleotide identity (ANI) and 0.33 alignment fraction (AF) are cut-offs set which are obtained by large scale genus comparisons (Auch, Jan, Klenk, & Göker, 2010; Barco et al., 2020; Richter & Rosselló-Móra, 2009). Further, criteria such as average amino acid identity (AAI) and percentage of conserved protein (POCP) have been proposed for genus delineation with 60-80% and 50% cut-offs, respectively (Konstantinidis & Tiedje, 2005; Qin et al., 2014). POCP is affected by genome size and hence cannot be applied for *Xylella* that has undergone extreme genome reduction (Hayashi Sant’Anna et al., 2019; Qin et al., 2014). However, core genome-based phylogenetic trees and AAI are not affected by genome reduction and can be employed to establish the identity and relationship of genera irrespective of genome size and/or GC content (Hayashi Sant’Anna et al., 2019; Indu, Ch, & Ch V, 2019; Konstantinidis & Tiedje, 2005). Interestingly, according to the genus demarcation values, minimum AF and ANI values for the XSXP phylogroup excluding *Xylella* were 0.35 and 76.54% respectively depicting *Xanthomonas, Stenotrophomonas* and *Pseudoxanthomonas* belong to same genus (supplementary table 1). Further, according to AAI cut-off value also all the strains of XSXP belong to the same genus i.e., *Xanthomonas* (figure 2a) and hence need to be merged into a single genus as *Xanthomonas.* Similarly, according to POCP cut-off of 50% all members of *Xanthomonas, Stenotrophomonas* and *Pseudoxanthomonas* belong to a single genus (figure 2b). Furthermore, within XX-MSG, 27 species formed clade I while other 6 species formed clade III. Further, ANI values clearly depicted that all the members of clade I and clade III belong to different species except for *X. alfalfa, X. perforans, X. euvesicatoria* and *X. gardneri, X. cynarae*, which represent heterotypic synonyms (supplementary figure 2). Clade II composed of *Xylella* genus. In case of S-MSG, the *Stenotrophomonas maltophilia* complex (Smc) consist of 6 species with two of them being clinical origin i.e. *S. maltophilia* and *S. africana*.

**Figure 2:**
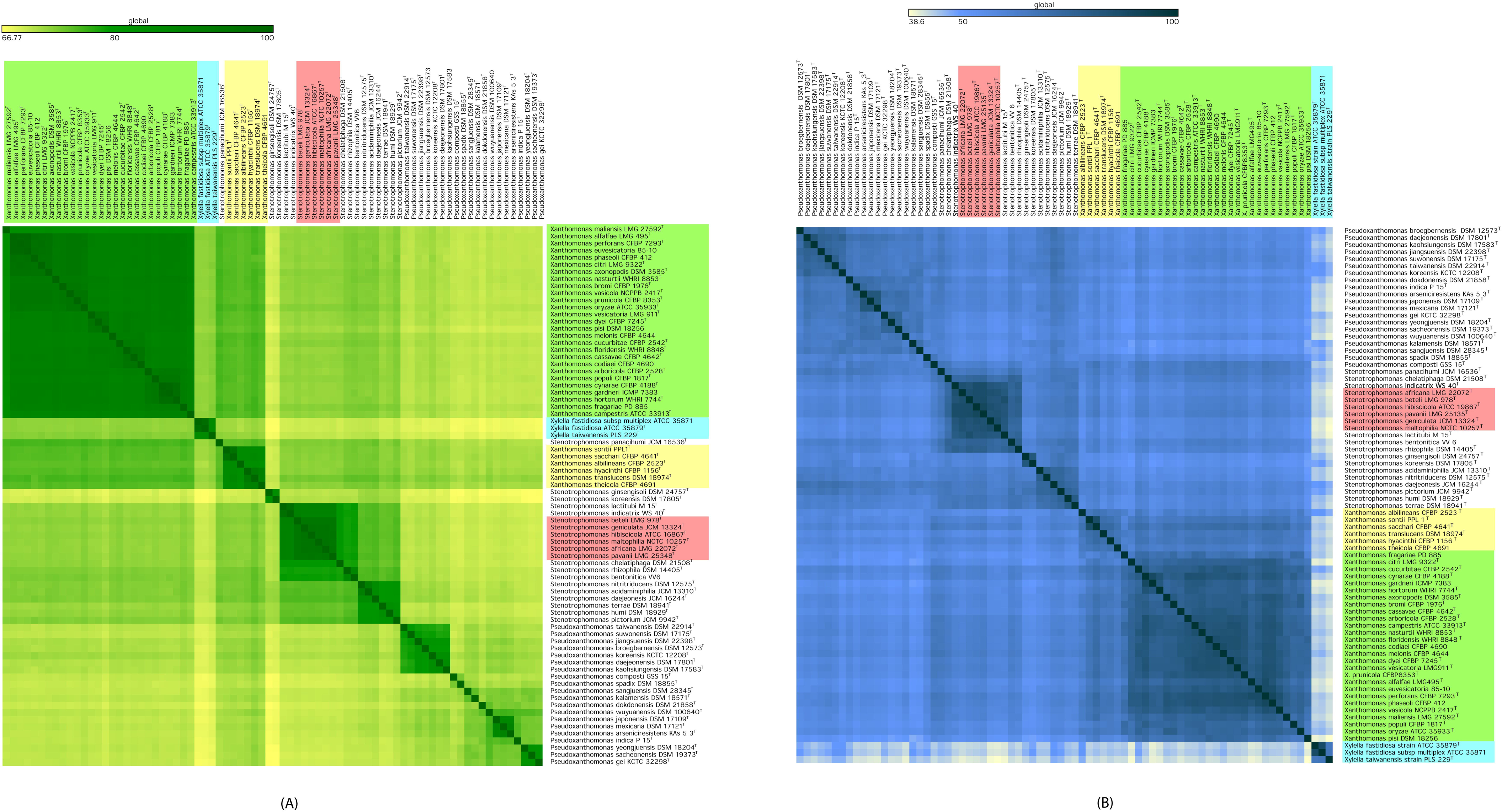
Heatmap of genome similarity for XSXP genomes. **a.** Matrix showing average amino acid identity (AAI) values amongst XSXP phylogroup. All the species complexes are represented in the colored boxes green: clade I; blue: clade II; yellow: clade III and red: clade IV. **b.** Heatmap showing percentage of conserved proteins (POCP) amongst XSXP group. All the species complexes are represented in the colored boxes green: clade I; blue: clade II; yellow: clade III and red: clade IV.

### Pan genome analysis and genome dynamics in XSXP

We performed pan genome analysis using roary v3.12.0 (Page et al., 2015) with 50% or greater amino acid identity revealed 47,624 gene families and 397 core genes for the XSXP phylogroup. Further, 48, 53, 205 and 175 genes were unique to clade I, III, II and IV respectively (supplementary table 2, 3, 4 and 5). Interestingly, clades II and IV constituting *Xylella* and Smc respectively are having higher number of unique genes. Inspection of clade IV unique genes revealed a type II secretion system, peroxidase, peptidases, efflux pumps and transporters like antibiotics/antimicrobials, fluoride ions, solvent, TonB dependent receptors, transcriptional regulators etc. (supplementary table 4). Further, unique genes of clade II belong to adhesin like type IV pili formation apart from glycosyltransferases, methyltransferases etc. (supplementary table 3). Overall COG classification revealed that hypothetical proteins are dominant class in all the clades suggesting unknown functions playing role in their success. Interestingly, in both clade I and III, COG class related to “signal transduction mechanisms” is second major class after hypothetical proteins, clade II is having more of “cell wall/membrane/envelope biogenesis”, “secondary metabolites biosynthesis, transport and catabolism” and “coenzyme transport and metabolism”. Whereas, clade IV was having more of “transcription”, “signal transduction mechanisms”, “intracellular trafficking, secretion and vesicular transport”, “cell wall/membrane/envelope biogenesis” and “inorganic ion transport and metabolism” (figure 3).

**Figure 3:**
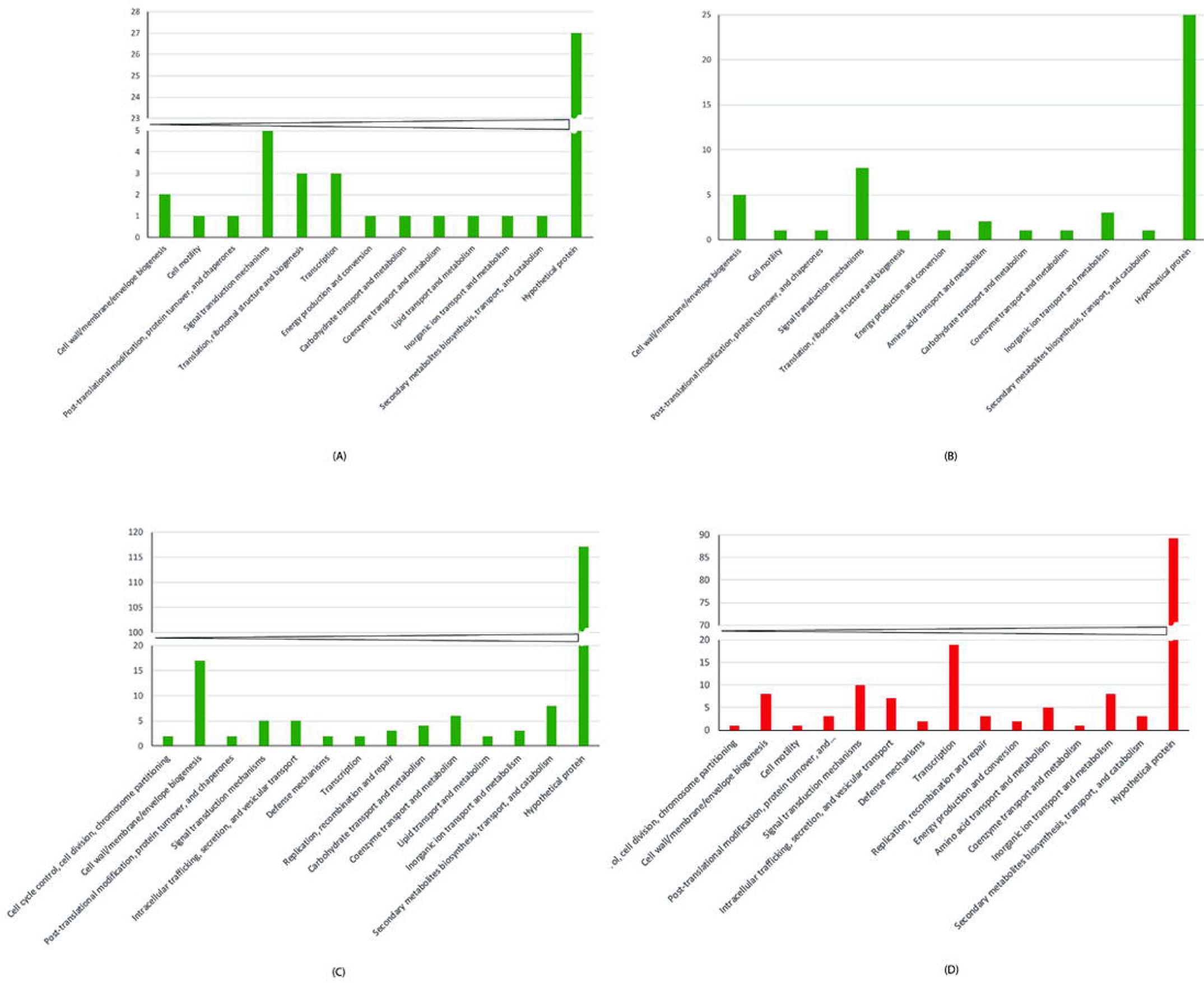
Distribution of COG-based functional categories of unique genes across several group. (A) clade I, (B) clade III, (C) clade II and (D) clade IV*. Stenotrophomonas maltophilia* complex represents the gene count across the several COG classes.

### Variable patterns of recombination within XSXP

To characterize genome-wide mosaicism in XSXP phylogroup we ran fastGEAR (Mostowy et al., 2017) on individual sequence alignments of the core and accessory genes or the pan genes. We found that 5,434 genes out of 47,162 pan genes had experienced recombination representing 11.5% of the pangenome (supplementary table 6). Here, 53,125 were the total recombining events detected out of which only 30% were detected from XX-MSG and remaining 70% were detected in genera *Pseudoxanthomonas* and *Stenotrophomonas* (figure 4). Amongst XX-MSG, least recombination events (72 out of 53125) were detected in *Xylella.* We observed heterogeneity in recombination sizes, majority of the recombination (~82%) were of less than 100bp recombining fragments while maximum recombining fragment was of 4469bp (supplementary table 6, 8). Further, 14 highway pairs were detected in the XSXP phylogroup, majority related to *Pseudoxanthomonas* and *Stenotrophomonas* (figure 5, supplementary table 7). Here, highways likely to represent specific lineages that function as hubs of gene flow. Here, highest recombination events were detected in *Pseudoxanthomonas* while least in *Xylella* strains.

**Figure 4:**
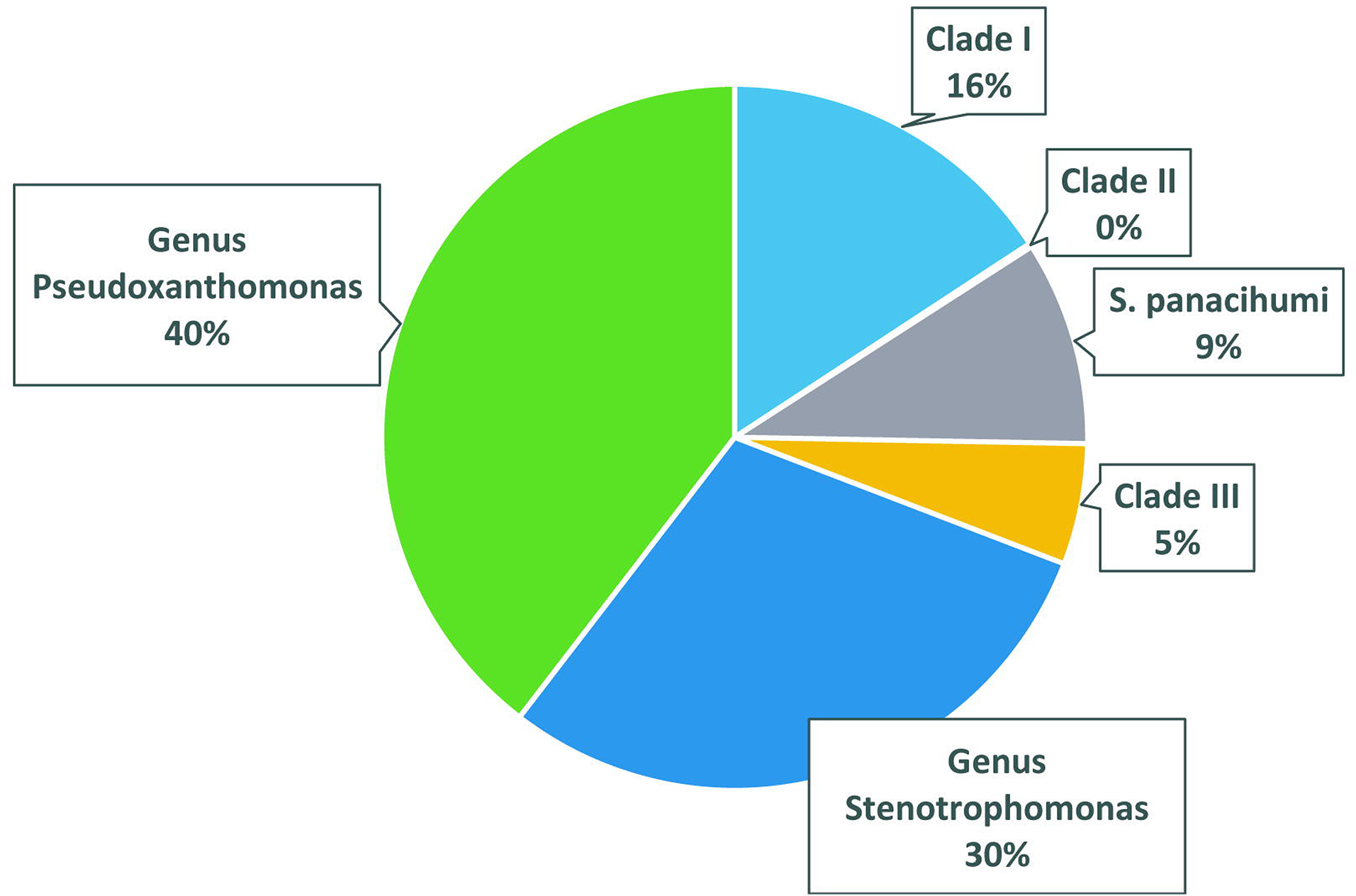
Number of recombination events detected in XSXP phylogroup.

**Figure 5:**
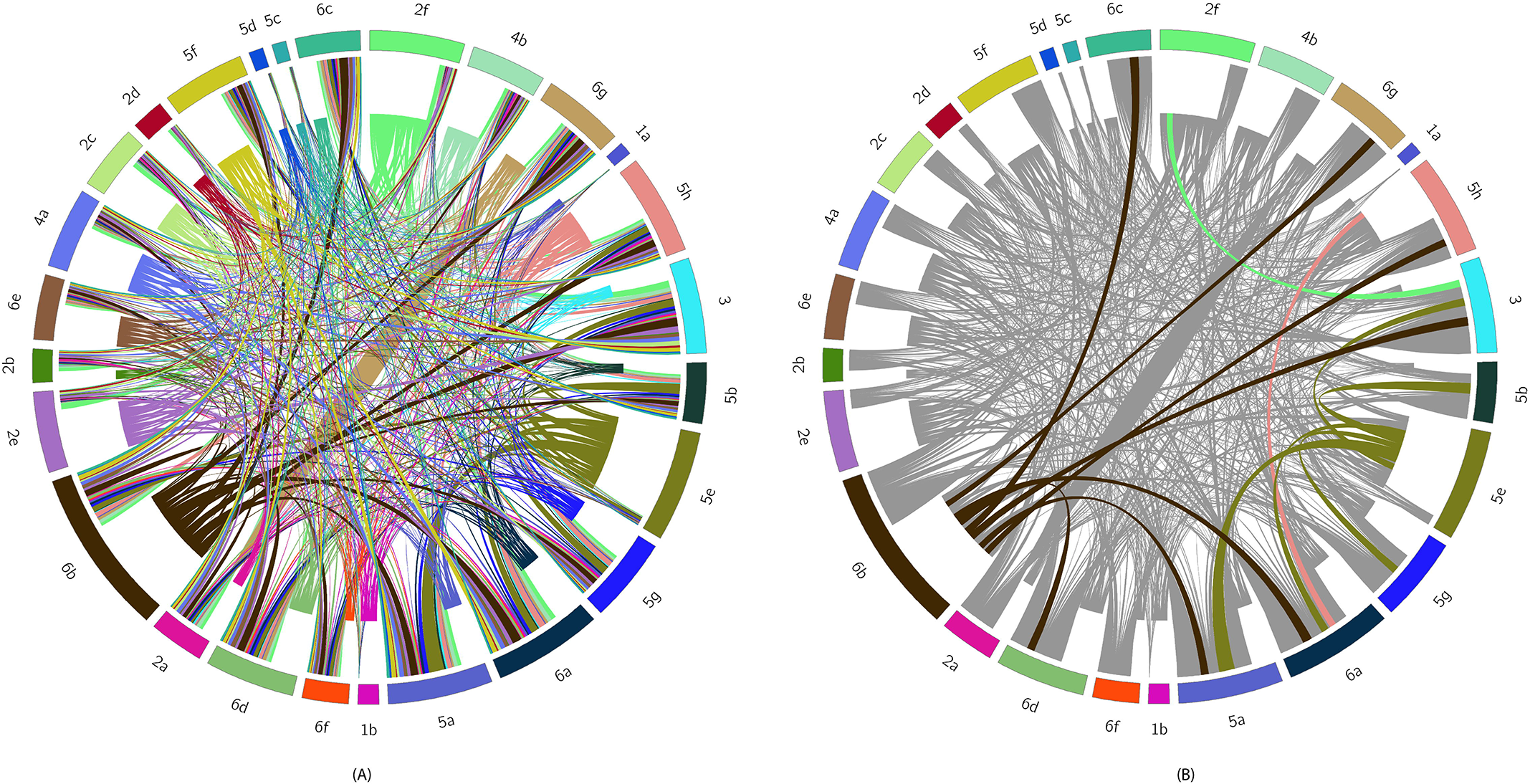
A) Recombination network generated by HERO. Outer ring represents sequence clusters and length of each fragment is proportional to number of recombination events affecting the SC. Ribbons connecting clusters that share recombination events and its thickness is proportional to the number of shared events. Color of the ribbon matches the donor cluster. B) Recombination network highlighting highways of recombination and non-highways are colored grey.

### Xanthomonadin pigment and xanthan exopolysaccharide gene cluster

Distinct yellow xanthomonadin pigment, responsible for yellow-colored colonies, encoded by *pig* gene cluster, and the thick mucus exopolysaccharide or xanthan encoded by *gum* gene cluster are characteristic features of canonical plant associated *Xanthomonas* species (He et al., 2020; Katzen et al., 1998; Rajagopal et al., 1997). Since, we are expanding breadth of *Xanthomonas* genus on the basis of phylo-taxono-genomics parameters, we scanned for the presence of these clusters in the genomes of XSXP constituent genera and species members (figure 6). Among the XX-MSG, the pig and gum gene clusters are present in all the members of *Xanthomonas* and *Xylella*, except for *X. theicola* and *X. albilineans* which are not having gum gene cluster. While both the clusters are highly conserved in sequence and distribution in clade I and II members of clade II show incomplete and possibly degenerated clusters, with *gumN, gumM, gumI, gumG* and *gmuF* absent from the *gum* cluster and orfs 8, 11, 13 and 14 absent from the pigment cluster.

**Figure 6:**
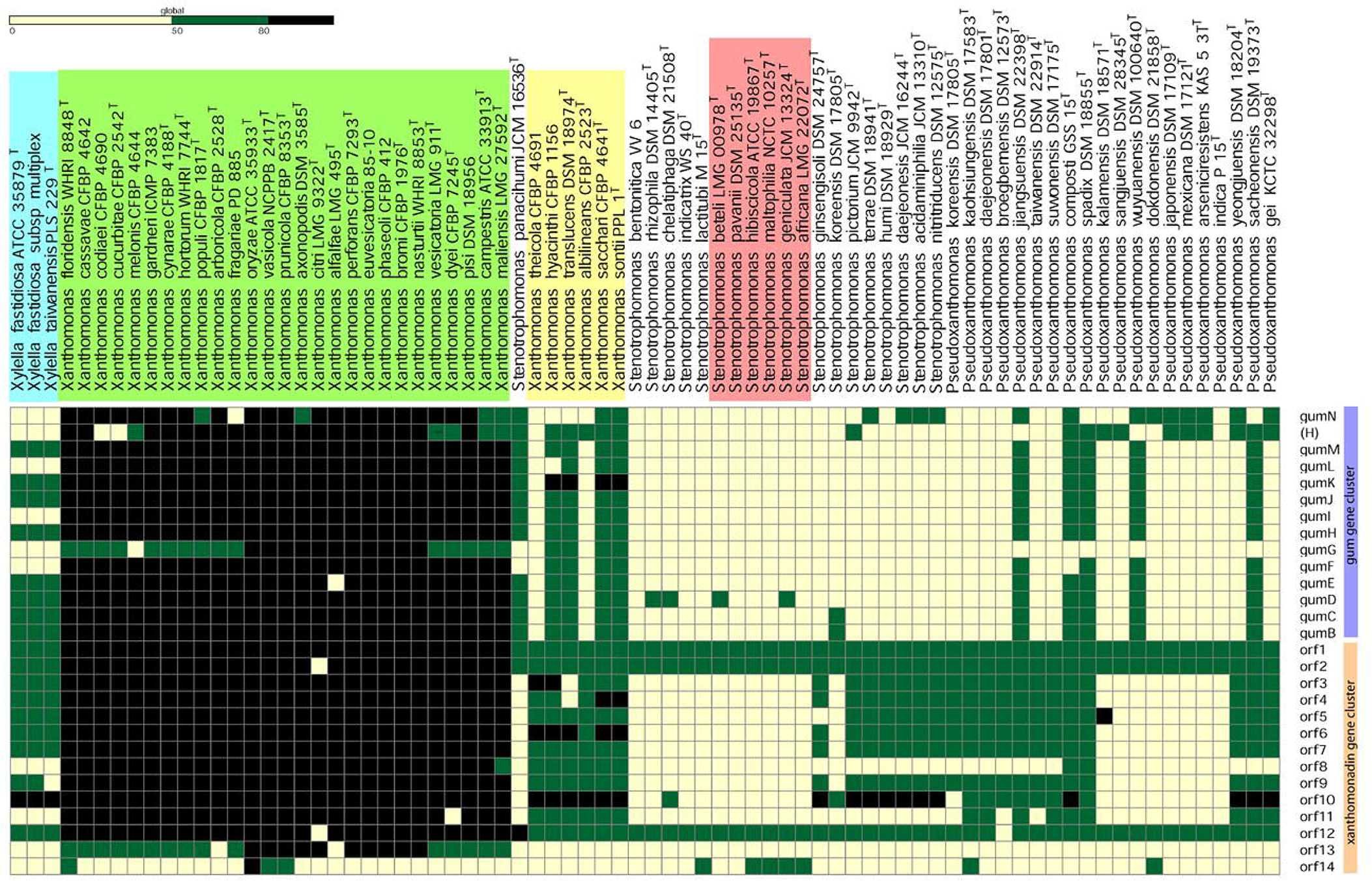
Heatmap showing status of *gum* gene cluster (upper panel) and xanthomonadin pigment (lower panel) in XSXP phylogroup. Here absence of the gene is denoted by yellow box and presence by black or green box. All the species complexes are represented in the colored boxes green: Clade I; blue: clade II; yellow: clade III and red: clade IV.

In members of other two genera understudy (*Stenotrophomonas* and *Pseudoxanthomonas*), these clusters are either absent or incomplete in majority of the members. Specifically, while all members of genus *Stenotrophomonas* lacks *gum* gene cluster, and 5 out of 20 species of *Pseudoxanthomonas* i.e., *P. jiangsuensis, P. composti, P. spadix, P. wuyuanensis* and *P. sacheonensis* harbour *gum* gene cluster. Whereas, xanthomonadin cluster is widely present in other two genera i.e., 7 out of 19 *Stenotrophomonas* and 13 out of 20 *Pseudoxanthomonas* species (figure 6). Remarkably, none of the clade IV of some clinical isolates have both the clusters.

## Discussion

The ability to sequence the genomes of bacterial strains in a cost-effective and high-throughput manner is transforming the way we understand genetics, phylogeny, and taxonomy. Inferring the phylogeny based on limited and highly-conserved sequence information such as 16S rRNA and housekeeping genes can be misleading not only at the species level but also at higher taxonomic levels (Sangal et al., 2016). Genomic investigation is the robust way to establish an organism’s identity, biology, and ecology in the proper context. Genome-based comprehensive taxonomic studies at various levels, such as order *Methylococcales,* order *Bacillales,* genus *Borrelia, and* genus *Lactobacillus*, have resolved the relationships and provided a robust basis for reclassification (De Maayer, Aliyu, & Cowan, 2019; Margos et al., 2019; Orata, Meier-Kolthoff, Sauvageau, & Stein, 2018; Salvetti, Harris, Felis, & O’Toole, 2018; Zheng et al., 2020). Whole genome studies using representative reference strains and type species of genera allowed us to resolve misclassifications at the level of families in the order *Lysobacterales* (*Xanthomonadales)* (Kumar et al., 2019). Whole genome studies also allowed the resolution of misclassifications at the level of species and clones (Bansal, Kumar, & Patil, 2021; Bansal, Midha, Kumar, & Patil, 2017; Kumar et al., 2019). In the case of genus *Stenotrophomonas*, we reported a large and hidden *Stenotrophomonas maltophilia* complex (Smc) consisting of at least nine species (Prashant P Patil, Kumar, Midha, Gautam, & Patil, 2018). Smc is a dynamically evolving group of human opportunistic pathogens with extreme drug resistance (Gautam et al., 2021; Kumar et al., 2020). However, deeper inter-genera and intra-genera genome-based investigations at the phylogenetic and taxonomic levels are lacking for the XSXP phylogroup. Considering the importance of the members of the XSXP phylogroup for plant and human health apart from their biotechnological potential, we performed an in-depth investigation by incorporating all the representatives of four closely-related genera through modern genome-based criteria.

Such a comprehensive phylo-taxono-genomic study revealed the role of evolutionary and ecological diversity in the formation of clades that actually belong to the genus *Xanthomonas* but have been historically classified into distinct genera such as *Xylella*, *Stenotrophomonas,* and *Pseudoxanthomonas*. The basal, multiple, and diverse groups comprising species of the genus *Pseudoxanthomonas* suggest it to be ancestral to the other three genera. Hence, it is not surprising that this large genus is named as *Pseudoxanthomonas* (Finkmann et al., 2000). At the same time, the presence of major lineage comprising the member species of *Stenotrophomonas* and another consisting of *Xanthomonas-Xylella* species suggests further specialization based on habitats or ecology. This phenomenon is quite obvious in the canonical plant-associated and pathogenic *Xanthomonas-Xylella* species that form a distinct mega-group unlike the mega-species group(s) represented by member species of *Stenotrophomonas* and *Pseudoxanthomonas* whose members are highly versatile. Besides, this finding indicates the role of lifestyle in the diversification and formation of clades that were previously reflected as separate taxonomic units at the level of genera when using only limited genotypic and phenotypic data. More importantly, we were also able to establish the robust phylogenetic relationship among the constituent member genera and species in XSXP using multiple phylogenetic approaches. In our case, this was possible because of the phylogenetic foundation provided by the species of *Pseudoxanthomonas* for the other three genera, asserting the significance of the genomic resource of the former. This proves the power of deep phylogenetic studies covering all member species and closely related genera.

Establishing the position of *Xylella*, which has a fastidious nature and an extremely lowered GC content along with highly reduced genome size, is of critical importance. While it is obvious that at the taxonogenomic level all the four genera are not distinct but belong to one genus, the element of surprise is that the genus *Xylella* is more related to *Xanthomonas* than to *Stenotrophomonas*. This variant lineage was confirmed by the sandwiched phylogenomic position of *Xylella* within the XX-MSG. The event that led to a sudden decrease in the GC content by >10% and genome size by ~50% is difficult to discern, and further studies in this regard are required. Our DEEPT studies imply that such an event could have happened during the diversification of *Xanthomonas* into pathogen of dicots from monocots or during diversification of *Xanthomonas* into a pathogen of monocots and dicots.

Conservation of the xanthomonadin pigment and *gum* gene clusters in clade I and clade III suggest to the essentiality of the pigment and the exopolysaccharide in the obligate plant-associated lifestyle. Since plants are directly and continuously exposed to light, it is important for a successful phytopathogen to have a unique pigment such as xanthomonadin (Rajagopal et al., 1997). It has been shown that xanthomonadin provides protection against photodamage in *X. oryzae* pv. oryzae that infects rice (Rajagopal et al., 1997). As plants are also known to produce large amounts of antimicrobial compounds and secondary metabolites (Ramírez-Gómez, Jiménez-García, Campos, & Campos, 2019), there is a need for a unique and thick polysaccharide coat. In fact, highly mucoid and distinct yellow-coloured colonies support our hypothesis that these clusters are evolved in the canonical *Xanthomonas* species from XX-MSG. Both these clusters were found to be more diversified in clade III than in clade I, which can be linked to the majorly dicot-associated lifestyle of the latter as compared to the majorly monocot associated lifestyle of the former. The leaves of monocots and dicots are different, which may affect the penetration of light and its effect on bacteria. Nonetheless, this hypothesis needs to be proven by further studies. Both the clusters were degenerated or incomplete in case of clade II, which can be associated with the dual lifestyle of *Xylella* that occurs within the foregut of the insect and within the plant where it is directly injected into the xylem (S. Chatterjee, R. P. P. Almeida, & S. Lindow, 2008b). Since *Xylella* is never exposed to light and plant defence response unlike the canonical *Xanthomonas* (i.e. clade I and III), it is not surprising that its clusters are not under natural selection. Genetic studies have revealed that the *pig* cluster is not required for virulence in plants (Rajagopal et al., 1997). Hence, the conservation of this cluster in clade I and III in the absence of its role in virulence suggests that the epiphytic mode is also crucial in plant adaptation (Pandey & Sonti, 2010).

The distributed presence of *pig* and *gum* clusters in a few members of XSXP outside the XX-MSG suggests that these clusters are ancestral and were present as incomplete clusters in the population even before the emergence of *Xanthomonas* clades. Eventually, they must have undergone diversification and selection when a member of XSXP came in contact with the plants. The ancestor of XX-MSG probably possessed both the clusters, thereby conferring an advantage for the plant-associated lifestyle. However, the presence of a primitive, ancestral, and incomplete pigment and *gum* gene clusters points that these clusters are not of foreign origin but were inherent to the ancestral XSXP population.

Even though *Xylella* seems to have undergone drastic genome reduction, the acquisition of a large number of unique genes with particular functions has played a key role in its emergence as a successful phytopathogen with insect/plant dual lifestyle. In both these variant lineages i.e. clade II and IV, the gaining of functions related to regulation suggest that apart from gene gain, the regulation of core or novel genes has also been responsible for their success. The commonality of the unique genes with regard to functions such as “intracellular trafficking, secretion, and vesicular transport”, “defence mechanism”, “replication, recombination, and repair” and “cell cycle control, cell division, and chromosome partitioning” reiterate their opportunistic, variant origin and parallels in evolution for dual/opportunistic lifestyle. Even though both exhibit dual lifestyle, unique gene analysis allowed us to pinpoint the functions associated with their success, such as adhesion in *Xylella* and efflux proteins/peroxidise (for multidrug resistance, adaptation to hospital setting, etc.) along with a novel type II secretion system in Smc. In an opportunistic human pathogen, a novel type II secretion system may compensate for the absence of type III secretion system in Smc (Crossman et al., 2008).

Even within the *Stenotrophomonas* group or S-MSG, the formation of a species complex corresponding to the MDR nosocomial pathogen points to the ongoing evolution and diversification as witnessed in the case of *Xylella* within XX-MSG. This finding is valuable in furthering our understanding of this emerging opportunistic human pathogen. Unique genes in clade I and III may be related to their adaptation to dicots and monocots, respectively. While many unique genes encode hypothetical proteins suggesting the importance of further functional genetic studies in clade I, II, III and IV, other major classes also provide clues regarding their evolution. One such hint is the importance of signal transduction in clade I and III, which can then serve as key targets for both basic and applied studies of dicot and monocot pathogens. Further systematic gene content analysis by excluding the variants within this group will also allow us to obtain insights into those genes that are important for the adaptation of *Xanthomonas* to monocots and dicots. Overall, our study reiterates the power and potential of systematic and large scale or deep taxonogenomics.

### Reconciliation, reconstitution and emended description of the genus *Xanthomonas*

Taxonomy in the pre-genomic era was dominated by classical polyphasic studies like phenotypic, microscopic, biochemical, fatty acids, quinines, etc. Any slight variation in the phenotypic data had led to assignment of new genus amongst *Xanthomonas* like species such as *Pseudomonas* (Hugh & Ryschenkow, 1961; Swings et al., 1983)*, Stenotrophomonas (Palleroni & Bradbury, 1993), Pseudoxanthomonas* (Finkmann et al., 2000) and *Xylella* (Wells et al., 1987)*. Pseudoxanthomonas* shares phenotypic traits with *Xanthomonas* like species such as colony color, cell shape, Gram staining, presence of branched chain fatty acid pattern and ubiquinone with eight isoprenoid units. Yet, *Xanthomonas* like strains with the ability to reduce nitrite but not nitrate to N_2_O and by the lack of C13: 0 iso 3-OH fatty acid were designated as genus *Pseudoxanthomonas* (Finkmann et al., 2000; Yang, Vauterin, Vancanneyt, Swings, & Kersters, 1993). Similarly, xylem-limited *Xanthomonas* like fastidious strains with GC content 51 to 53 mol % were designated as separate genus *Xylella* (Wells et al., 1987). Presence of fimbrae, multiple flagella and inability to produce xanthan gum and xanthomonadins distinguished another new genus *Stenotrophomonas* amongst the *Xanthomonas* like species (Palleroni & Bradbury, 1993).

Even then, these genera have characteristic common presence of branched chain fatty acid pattern and an ubiquinone with eight isoprenoid units (Q-8) (Finkmann et al., 2000; Swings et al., 1983). Hence, classification of *Xanthomonas* like species as distinct genera has been controversial which led to several reclassifications (Lee et al., 2008; Palleroni & Bradbury, 1993). Also plant or human associated microbes have influenced their assignment into separate genera. Advent of genomics have highlighted role of horizontal gene transfer in diverse lifestyles of these microbes. Acquisition of genes would have provided much needed functions in adaption to humans (in case of *S. maltophilia* complex) (Prashant P. Patil et al., 2021) and insect host (in case of *Xylella*) (Chatterjee et al., 2008b).

Hence, there is a need to reconcile the published phenotypic data in the light of genomic data and relationship of all the four genera. The current standing in nomenclature of family *Lysobacteraceae* is based on CSI phylogeny (using only 28 conserved proteins) which describes these *Xanthomonas* like species into distinct genera namely *Xanthomonas, Xylella, Stenotrophomonas* and *Pseudoxanthomonas* (H. S. Naushad & Gupta, 2013; S. Naushad et al., 2015). Although CSI phylogeny itself suggests taxonomic position of *Xylella* in between *Xanthomonas* group 1 and group 2 which is not a true evidence for considering *Xanthomonas* and *Xylella* as distinct genera (S. Naushad et al., 2015). Hence, there is a need to consider them as synonyms of genus *Xanthomonas*.

In the present study we provide whole genome based deep-phylo-taxonogenomic evidences which clearly depicts the existence of unary genus for all *Xanthomonas* like species (*Xanthomonas, Xylella, Stenotrophomonas* and *Pseudoxanthomonas*). Here, we have implemented three different robust approaches to obtain core-genome phylogeny of XSXP phylogroup i.e. PhyloPhlAn (>400 genes), roary (382 across 99-100% isolates) and PIRATE (1149 genes across 95-100% isolates). Core genome phylogeny obtained from these methods reveals sandwiched positioning of *Xylella* between two groups of *Xanthomonas.* While, *Pseudoxanthomonas* was ancestral among XSXP phylogroup. Further, genus delineation using ANI, AF, AAI, POCP clearly depicted *Xanthomoans, Stenotrophomonas* and *Pseudoxanthomonas* as single genus. However, *Xylella* cannot be assessed using genome similarity assessment due to its drastic reduction in genome size (leading to low number of CDS and GC content). In light of deep phylo-taxono-genomics findings along with published polyphasic data XSXP phylogroup warrants reunification of all its *Xanthomonas* like species.

## Materials and Methods

### Genome procurement from public repository

A total of 76 strains were used in the present study. Genome sequences of 33, 20, 20 and 3 were type strains of *Xanthomonas, Stenotrophomonas* and *Xylella*, were available from the NCBI database (https://www.ncbi.nlm.nih.gov/). All the genomes were then accessed using checkM v1.1.0 (Parks, Imelfort, Skennerton, Hugenholtz, & Tyson, 2015) with the cutoff of less than 3 percent contamination and completeness (table 1).

### Whole genome-based phylogeny

Phylogenomic analysis based on more than 400 putative conservative genes was carried out using PhyloPhlAn v3.0 (Asnicar et al., 2020). Here, USEARCH v5.2.32 (Edgar, 2010), MUSCLE v3.8.3 (Edgar, 2004) and FastTree v2.1 (Price, Dehal, & Arkin, 2010) were utilized for orthologs searching, multiple sequence alignment and phylogenetic construction respectively. All strains of XSXP phylogroup and *Luteimonas mephitis* DSM12574T (used as an outgroup) were used for construction of the phylogeny.

In order to obtain a more robust whole genome phylogeny, core genome based phylogeny was constructed using MAFFT v7.31 (Nakamura, Yamada, Tomii, & Katoh, 2018) (https://mafft.cbrc.jp/alignment/software/) and the FastTree v2.1 (Price et al., 2010) which was integrated within the roary v 3.12.0 (Page et al., 2015) with identity cut-off of 60%. Implementation of the tools can be found in detail under the section of pan genome analysis.

As here, XSXP phylogroup consist of divergent genomes and to further confirm the core genomes deduced by roary and their phylogeny, we have used PIRATE (Bayliss et al., 2019). This is better known to identify orthologue groups of diverged gene sequences by using amino acid identity threshold of 50%, 60%, 70%, 80%, 90% and 95%.

### Taxonogenomic analysis

Genome relatedness was assessed using the average amino acid identity (AAI), percentage of conserved protein (POCP) and average nucleotide identity using (ANI). AAI was calculated using CompareM v0.0.23 (https://github.com/dparks1134/CompareM), which uses the mean amino acid identity of orthologous genes between a given pair of genomes. POCP is another method to evaluate the genome relatedness at genus level, which is based on amino acid conservation. POCP is calculated with the blast search using the default settings (Qin et al., 2014) (https://figshare.com/articles/POCP-matrix_calculation_for_a_number_of_genomes/5602957). Further, FastANI v1.3 (Jain, Rodriguez-R, Phillippy, Konstantinidis, & Aluru, 2018) an alignment-free sequence mapping method with default settings was used to calculate ANI values with default settings.

### Pan genome analysis

Pan genome analysis of the strains was performed using Roary v3.12.0 (Page et al., 2015). Here, gff files generated by Prokka v1.13.3 (Seemann, 2014) were used as input for Roary pan genome analysis. Since, we are analyzing genomes from different species, the identity cut-off used was 60%. Functional annotation of the core genes identified was performed using eggNOG v4.5.1 (Huerta-Cepas et al., 2015).

### Recombination analysis

To identify recombination, we used fastGEAR (Mostowy et al., 2017)with default parameters on individual genes of the pan genome identified by Roary. HERO uses output of fastGEAR to identify donors and recipients in the recombination events. It also mapped recombination events to the sequence clusters identified by core genome-based phylogeny (supplementary table 6) to elucidate potential drivers of biases in recombination partners.

The custom script has been provided with HERO on its GitHub page (https://github.com/therealcooperpark/hero) as “sidekick.py” for reproducibility and convenience when using HERO in similar workflows.

### Cluster analysis

Protein sequences of the gene clusters (Bansal, Midha, et al., 2019b) were used as query and tBLASTn searches were performed on the XSXP phylogroup genomes. Here, tBLASTn searches were performed using standalone BLAST+ v2.9.0 (Camacho et al., 2009) and cut-offs used for identity and coverage were 50% and 50% respectively. Heatmap for the blast searches were generated using GENE-E v3.03215 (https://software.broadinstitute.org/GENE-E/).

## Author Contributions

KB and SK carried out all the bioinformatics analysis. KB, SK and PBP drafted the manuscript with inputs from AK and AS. PBP has conceived the study and participated in the design. All the authors have read and approved the manuscripts.

## Acknowledgement

This work was supported by a project entitled “Megagenomic and Metagenomic insights into adaptation and evolution of fruit Microbiome” GAP0187 of CSIR to P.B.P.

## Figures and Tables legends

**Supplementary figure 1:** Core genome phylogeny of the XSXP phylogroup. Here, core genome alignment is obtained by using PIRATE.

**Supplementary figure 2:** OrthoANI value calculated for the XSXP phylogroup.

**Supplementary table 1**: Average nucleotide identity (ANI) and alignment fraction (AF) amongst the XSXP phylogroup.

**Supplementary table 2**: Genes unique to clade I, its COG classification and its protein sequence.

**Supplementary table 3**: Genes unique to clade III, its COG classification and its protein sequence.

**Supplementary table 4**: Genes unique to clade II, its COG classification and its protein sequence.

**Supplementary table 5**: Genes unique to clade IV, its COG classification and its protein sequence.

**Supplementary table 6**: Recombination events detected within XSXP phylogroup.

**Supplementary table 7**: Sequence clusters among the XSXP phylogroup.

**Supplementary table 8**: Genes undergoing recombination amongst the XSXP pangenome.

