## Supplementary figures and images for "Deep phylo-taxono genomics reveals *Xylella* as a variant lineage of plant associated *Xanthomonas* with *Stenotrophomonas* and *Pseudoxanthomonas* as misclassified relatives"

### Supplementary Figure 1

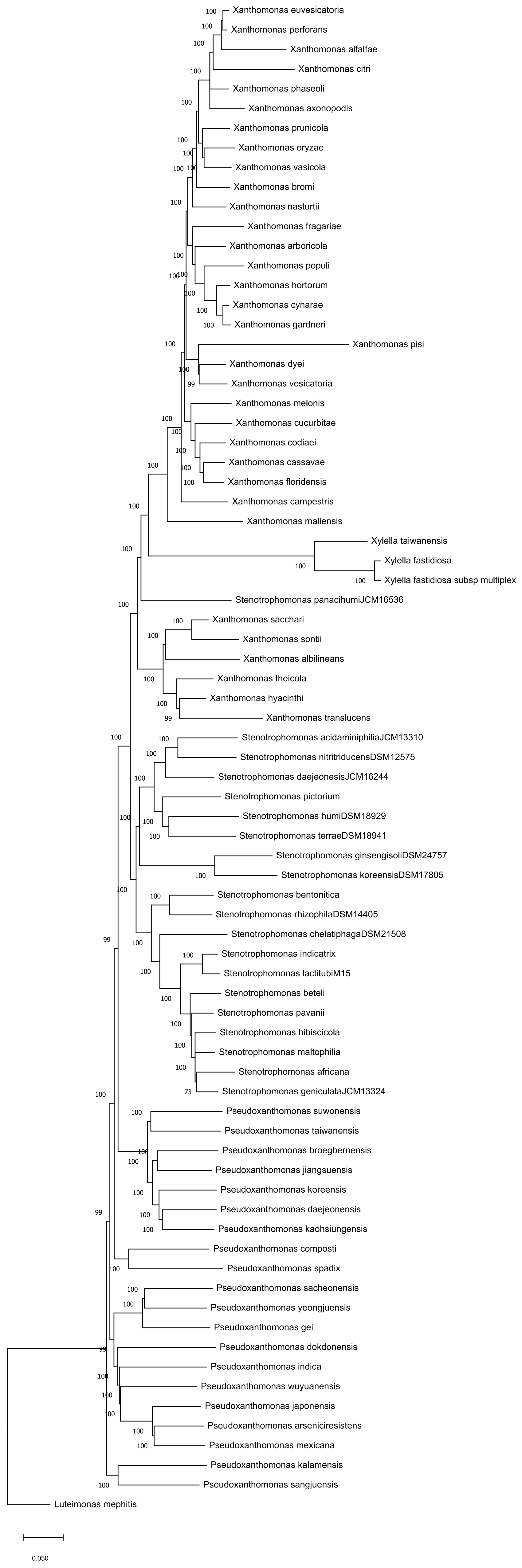

### Supplementary Figure 2

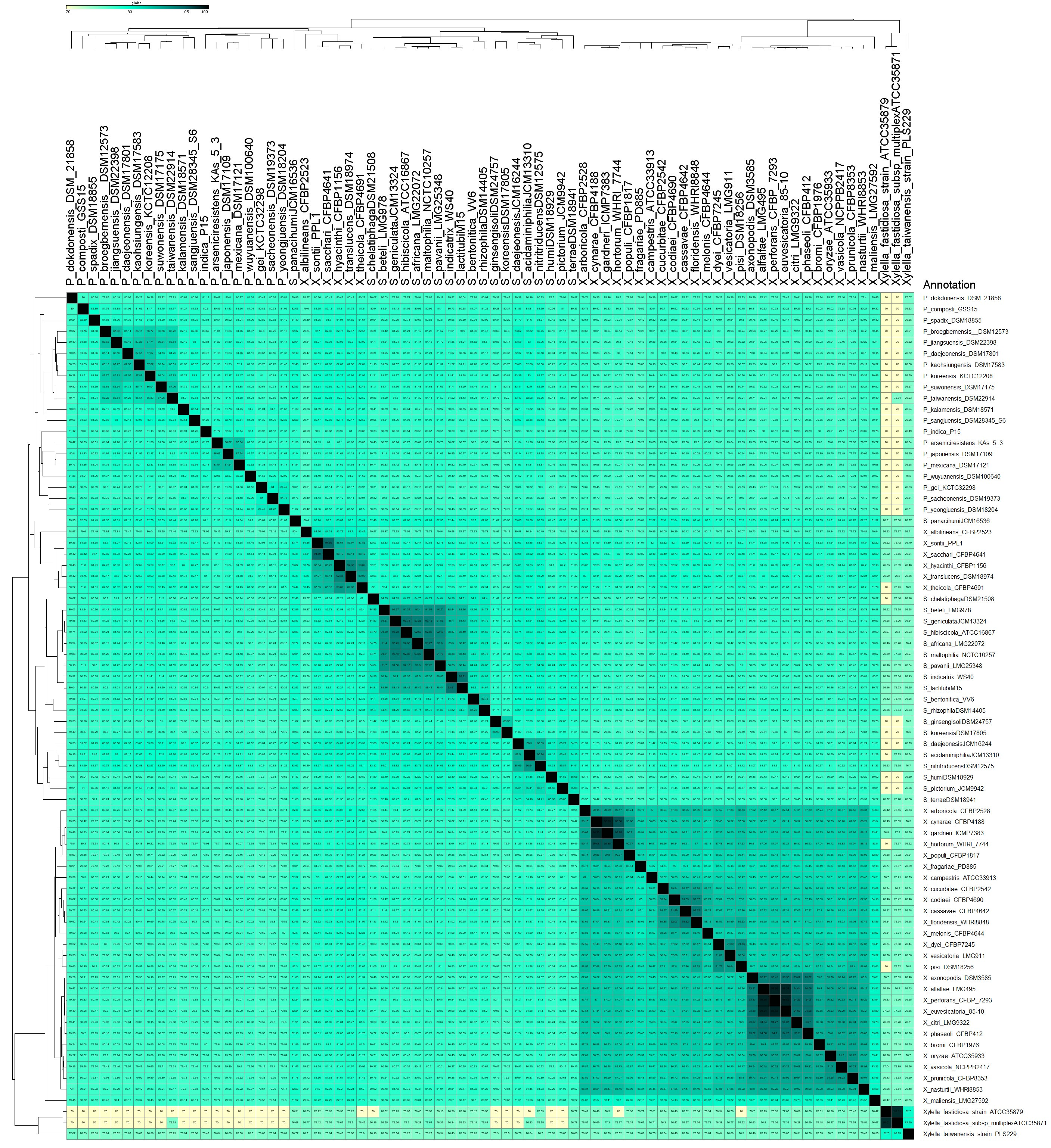
